# Single-cell Profiling of Airway-associated CD4^+^ T cells in Metal-induced Hypersensitivity

**DOI:** 10.64898/2026.11.11.765065

**Authors:** Jarren Ypil, Rui Fu, Masoom Raza, Li Li, Lisa Maier, Andrew P. Fontenot, Charles A. Dinarello, Shaikh M. Atif

## Abstract

Omics-based approaches are increasingly used to identify unique molecular signatures that may serve as biomarkers for early disease detection and potential therapeutic targets. Chronic beryllium disease (CBD), resulting from beryllium-induced hypersensitivity, is characterized by the accumulation and expansion of pathogenic CD4^+^ T cells in the lungs, ultimately contributing to impaired pulmonary function. In this study, we performed single-cell RNA sequencing (scRNA-seq) on sorted viable bronchoalveolar lavage (BAL) cells from beryllium-exposed patients (n=2) and compared their transcriptional profiles with publicly available scRNA-seq datasets from healthy controls (n=10). We identified 11 distinct CD3-associated cell clusters comprising T cells, natural killer (NK) cells, and regulatory T cells. Further analysis of CD4^+^ T cells from beryllium-exposed patients revealed increased expression of genes associated with T cell activation, immune regulation, and persistence, including *IFNG*, *TNF*, *TNFAIP3*, and *CD74*. Gene Ontology (GO) enrichment analysis further revealed upregulation of pathways associated with cellular activation, immune responses, and immune effector functions, whereas pathways related to metabolic processes were downregulated in beryllium samples compared with healthy controls. Overall, our findings identify distinct transcriptional signatures, including *IFI27* and *FABP4*, in CD4^+^ T cells from beryllium-exposed patients, providing insights into the molecular mechanisms underlying beryllium-induced pulmonary inflammation and highlighting potential candidates for further investigation as disease biomarkers and therapeutic targets.

## Introduction and Results

Chronic beryllium disease (CBD) is a metal-induced hypersensitivity disorder characterized by CD4^+^ T cell alveolitis (1–3). Single-cell gene expression profiling of bronchoalveolar lavage (BAL) cells focused on T cells provides insight into unique subsets of CD4^+^ T cells, functional pathways, and/or molecules that could potentially be developed into novel therapeutics for CBD and other granulomatous lung diseases.

To understand the unbiased cellular composition of airway-associated CD4^+^ T cells in subjects with beryllium exposure, we used BAL cells from beryllium (Be)-exposed subjects (n=2) diagnosed using previously defined criteria (4, 5) to comprise the disease cohort. The bronchoalveolar lavage samples were collected from Be-exposed subjects at National Jewish Health (NJH) Granuloma Clinic. Subjects provided informed consent before inclusion in the study, and the protocol was approved by the Human Subjects Institutional Review Boards at the University of Colorado Anschutz Medical Campus and National Jewish Health. We profiled 9118 BAL cells from these two Be-exposed subjects (CBD and BeS) and 10 healthy individuals whose datasets were publicly available (Fig. 1A). After performing quality control and normalizing the data, analysis of 2378 CD3 T cells revealed a total of 11 clusters among the disease and healthy cohorts based on expression of signature genes using the tool suite Seurat (6). Data were visualized using a uniform manifold approximation and projection (UMAP) plot (Fig. 1A-C) (7), and established markers were used for cluster annotation and gene identification (8). Using the gene signatures *CD4*, *CD8A*, *FOXP3*, *KLRC1,* and *KLRC3*, the 11 distinct CD3 clusters were composed of CD4, CD8, regulatory CD4 T cells (Tregs), NK cells, and a separate unknown cluster dominated by a mixed population of T cells expressing CD4 and CD8 (cluster 11), which was excluded from further analysis. Based on the closeness of the gene profiles for CD8^+^ and CD4^+^ T cells, clusters 7, 8, and 9 were combined into a single CD8^+^ T cell cluster, and clusters 1, 2, 3, 4, and 5 were merged into a CD4^+^ T cell cluster (Fig. 1C). The Treg cluster was kept separate based on its distinct location in the UMAP (Fig. 1C).

**Figure 1:**
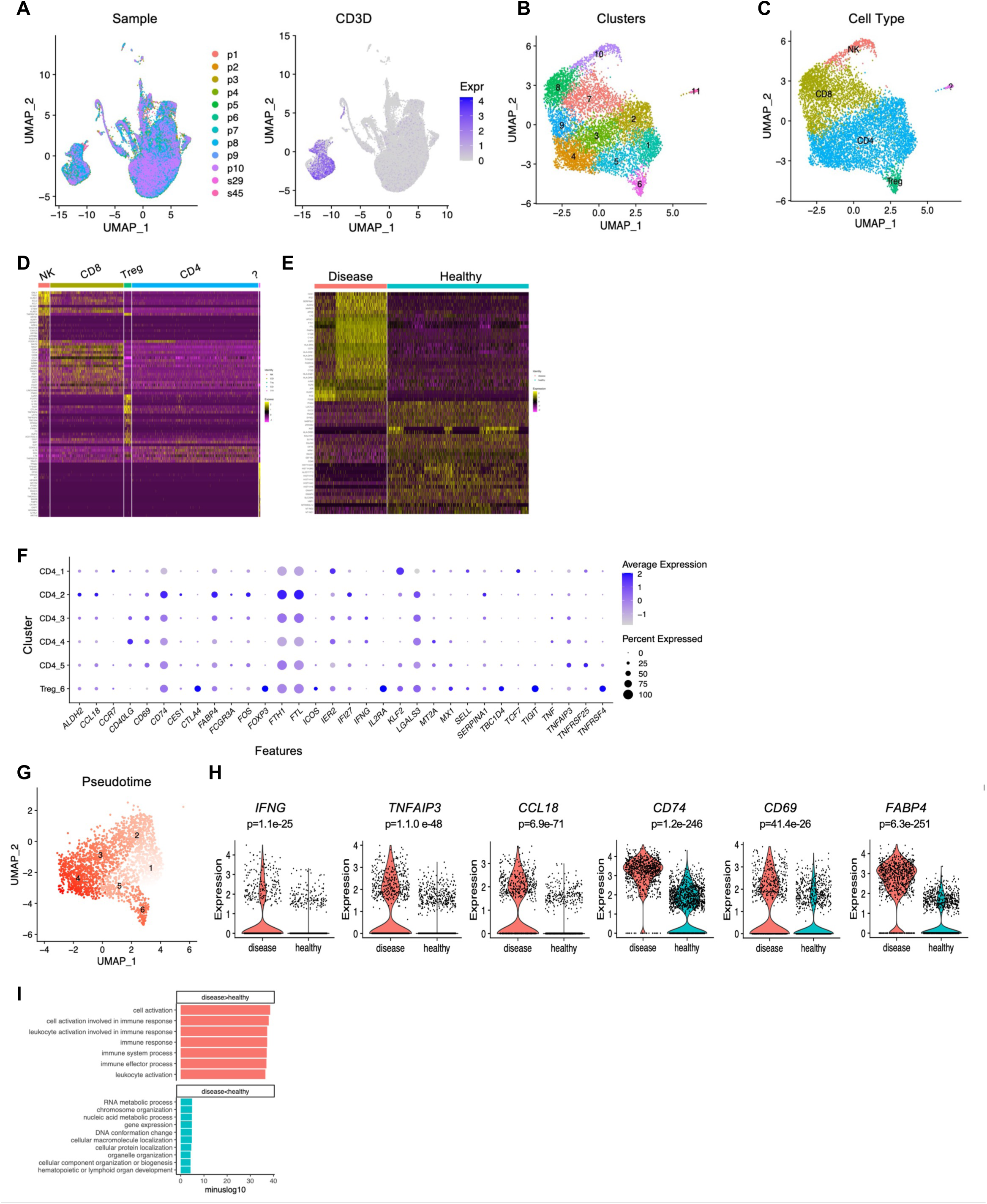
Single-cell transcriptomics of bronchoalveolar lavage cells show unique CD4^+^ T cell subsets encompassing cellular heterogeneity. A) UMAP plot of bronchoalveolar lavage (BAL) cells from disease and healthy control subjects shows cellular distribution (left) and the CD3 T cell cluster (right) present in all groups. B) Colored UMAP shows 11 clusters in the CD3 subset of cells. C) Combined clusters of cells based on normalized gene signatures. D) Heatmap shows differentially expressed genes within NK, CD8, Treg, and CD4^+^ T cell clusters. E) Heatmap shows CD4^+^ T cell transcriptome profile comparing disease and healthy individuals. F) Dot plots show variation in the expression of signature genes distinguishing various clusters of CD4^+^ T cells (1–6). G) Pseudotime trajectory shows that the acquisition of activation status of CD4^+^ T cells changes from cluster 1 (activation) to cluster 4 (effectors). H) Violin plots show expression levels of genes associated with effector (*IFNG, TNFAIP3, CD74,* and *CCL18*), activation, and regulation (*CD69 and FABP4*) of CD4^+^ T cells. I) Bar plots show minus log10 values from enrichment analysis of representative Gene Ontology (GO) Biological pathways in CD4^+^ T cells from disease and healthy controls. P-value < 0.01 was statistically significant.

The transcriptional profiles of clusters for both subject cohorts corresponding to NK cells, CD8^+^ T cells, CD4^+^ T cells, and Tregs are shown in Fig. 1D. However, since beryllium-induced disease is predominantly a CD4^+^ T cell-mediated disorder (3, 9), we focused on a detailed transcriptome analysis of CD4^+^ T cells, including Tregs. Differential expression analysis of CD4^+^ T cells was performed to compare gene profiles in Be-diseased individuals to those of healthy control subjects (Fig. 1E). Furthermore, we examined the percent expression of genes defining the functions of CD4^+^ T cells in different clusters of CD4^+^ T cells on a per-cell basis using dot plots (Fig. 1F). We found a large percentage of cells expressing T effector cytokines (*IFNG*, *TNF* and *TNFAIP3*) were localized in CD4 clusters 3 and 4, while CD74, an invariant chain molecule expressed on memory T cells and linked to inflammation (10), was dominantly expressed in CD4 cluster 2. We also observed increased expression (∼2-fold) of ferritin light chain and heavy chain, *FTL*, *FTH1*, and *IFI27* in cluster 2, distinguishing it from the other CD4 clusters. To date, there is no known direct role of ferritin in CD4^+^ T cell function, but *IFI27* gene expression is increased in viral and autoimmune diseases (11, 12). Like CD4 clusters 2, 3, and 4, clusters 1 and 5 were distinguished by higher cluster-specific expression of *IER2, KLF2,* and *TNFRSF25* than the other clusters, and cluster 6 (Tregs) showed high expression of *FOXP3, TIGIT,* and *CTLA4*, genes that were minimally expressed in other clusters. Overall, the dot plot data suggested that the CD4^+^ T cell clusters were interrelated and consisted of many overlapping gene sequences. To investigate relatedness among clusters, we performed a Pseudotime trajectory analysis (Fig. 1G) that identifies the association among the CD4^+^ T cell clusters (13). Based on the expression of *SELL*, *CCR7*, and *KLF2* in cluster 1 (Fig. 1F), we considered this cluster to be composed of recently migrated cell populations from which the continuum of cells occurs, giving rise to a terminally differentiated effector population in cluster 4, as evident by the expression of *IFNG*, *TNFAIP3*, and *CCL18* (Fig. 1F). These findings are consistent with our murine studies, in which RNA sequencing of purified lung CD44^+^CD4^+^ T cells from BeO-exposed HLA-DP2 Tg mice revealed a similar upregulation of genes (*Ifng*, *Tnf*, and *Cd74*) associated with T cell activation and function (14).

Next, we examined a subset of genes involved in CD4^+^ T cell activation and regulation among 249 differentially expressed genes (DEGs) between disease and healthy individuals (Table 1). DEGs were defined by a Wilcoxon signed-rank test, with thresholds of adjusted p-value ≤ 0.01 and log2 fold change of over 0.25. CD4^+^ T cells in the Be-exposed group showed significantly higher levels of effector and activation markers and cytokines, including *IFNG*, *TNFAIP3*, *CD69*, *CD74*, *CCL18*, and fatty-acid binding protein 4 (*FABP4)*, which supports the maintenance, longevity, and function of CD8^+^ T cells (10). The increased expression of this gene in a CD4^+^ T cell cluster suggests a role in the metabolism and function of CD4^+^ T cells (Fig. 1H). We also used GO pathway enrichment analysis to identify unique pathways upregulated in Be-exposed CD4^+^ T cells compared to healthy controls (Fig. 1I). The top 200 DEGs, ranked by adjusted p-value, were used for GO term enrichment analysis. Our data show upregulation of pathways involved in cell activation, immune responses, and immune effector processes, while pathways involved in metabolic processes, protein localization, and transcriptional regulation of genes were downregulated in disease subjects relative to healthy controls.

**Table 1.**
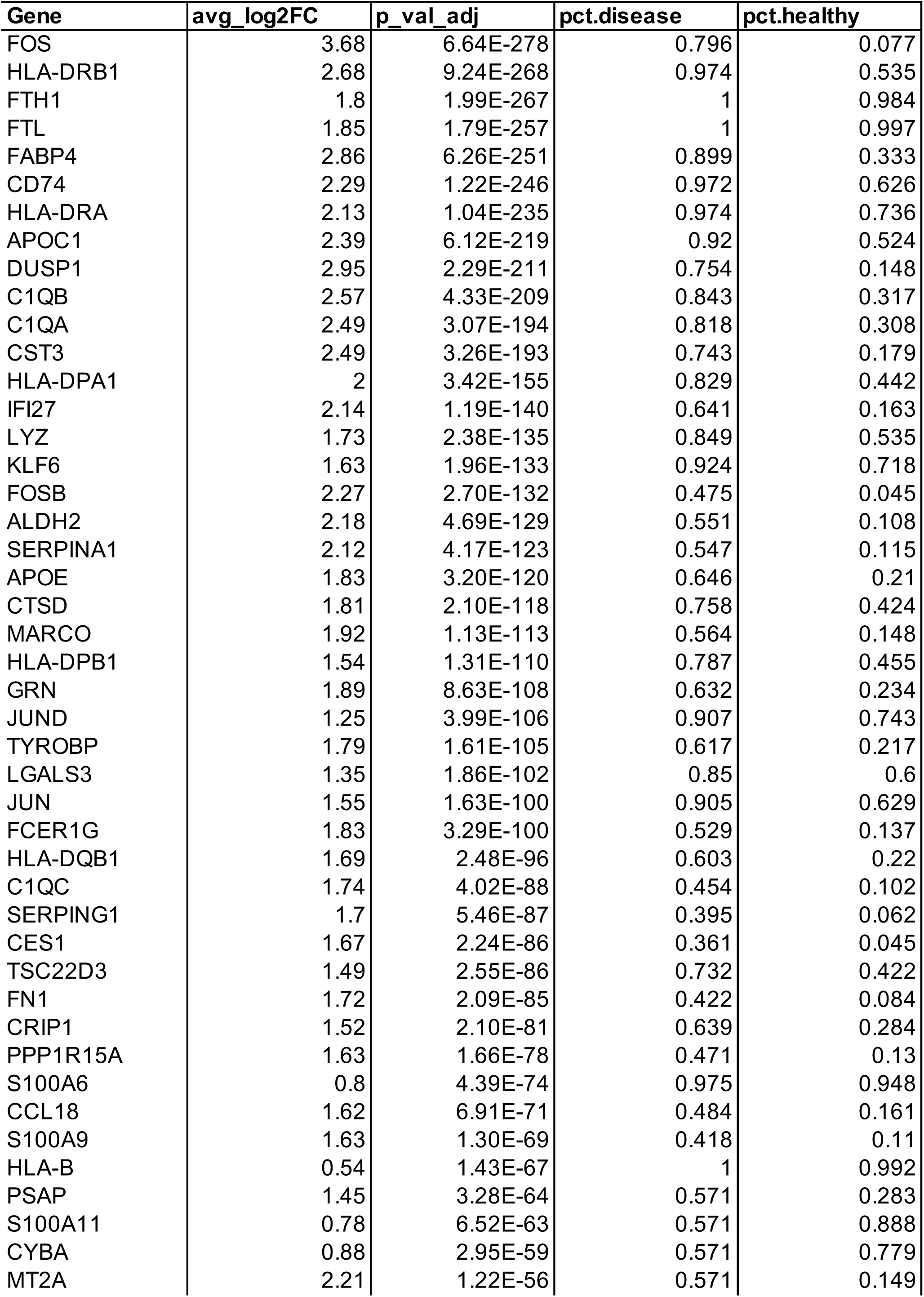
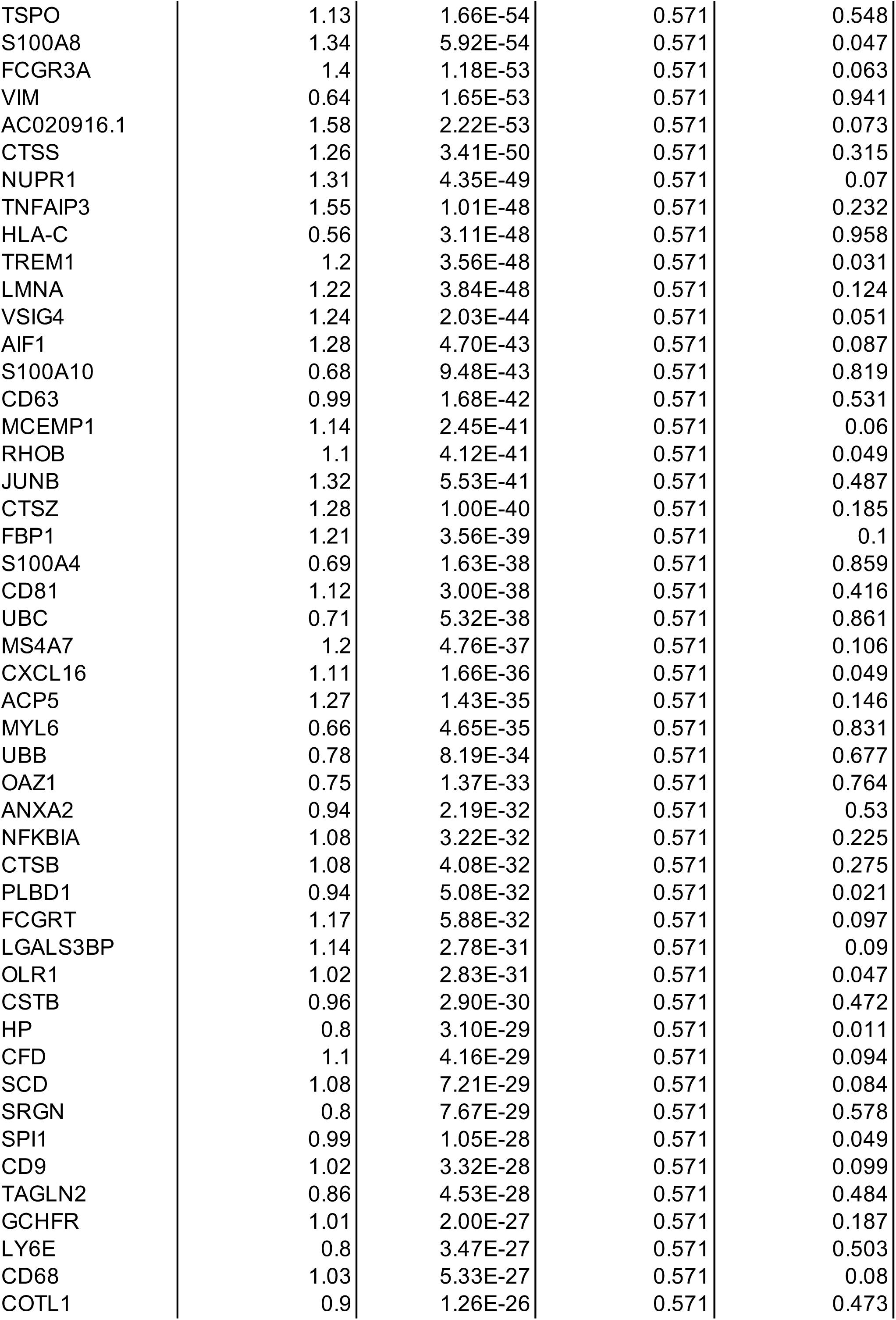
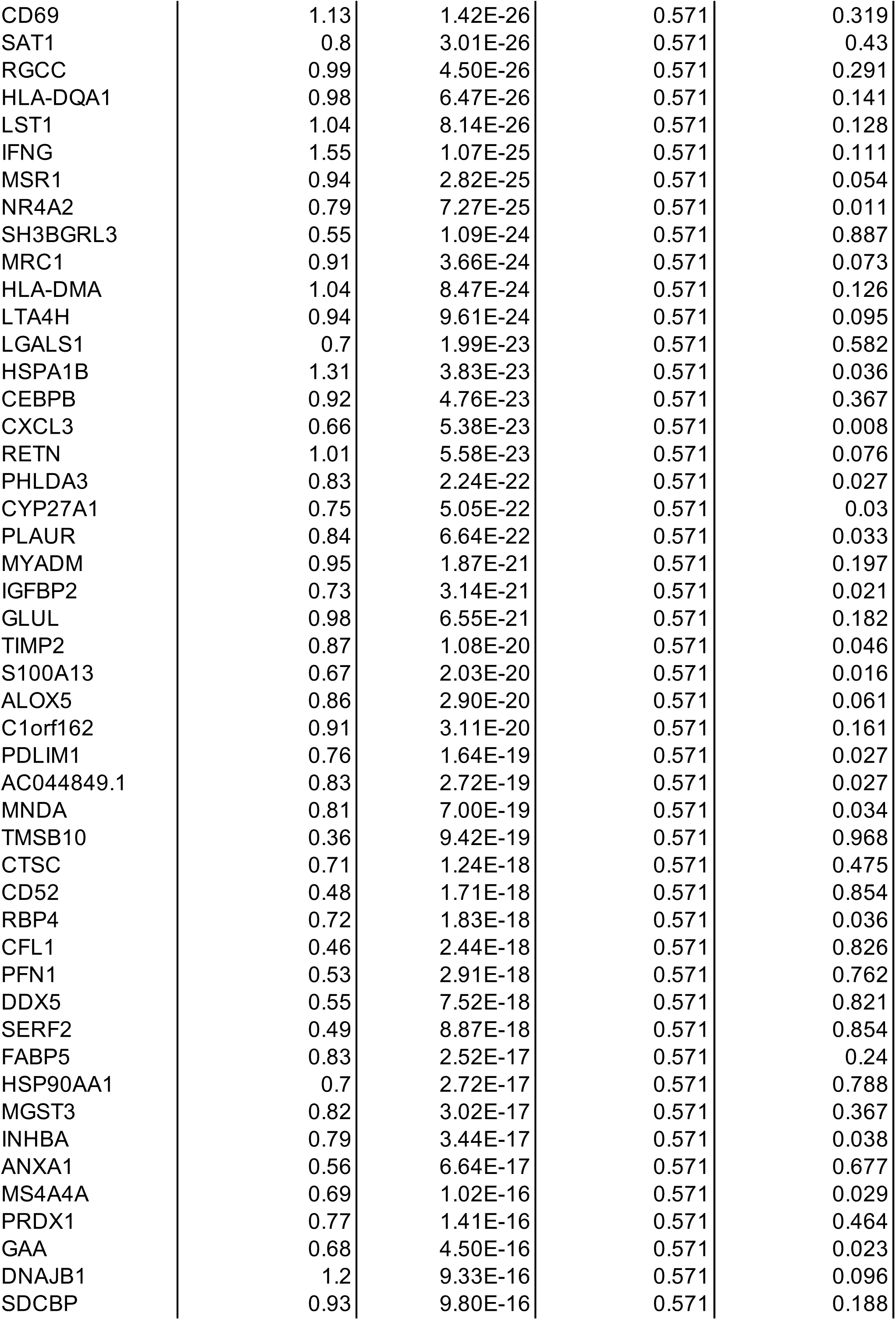
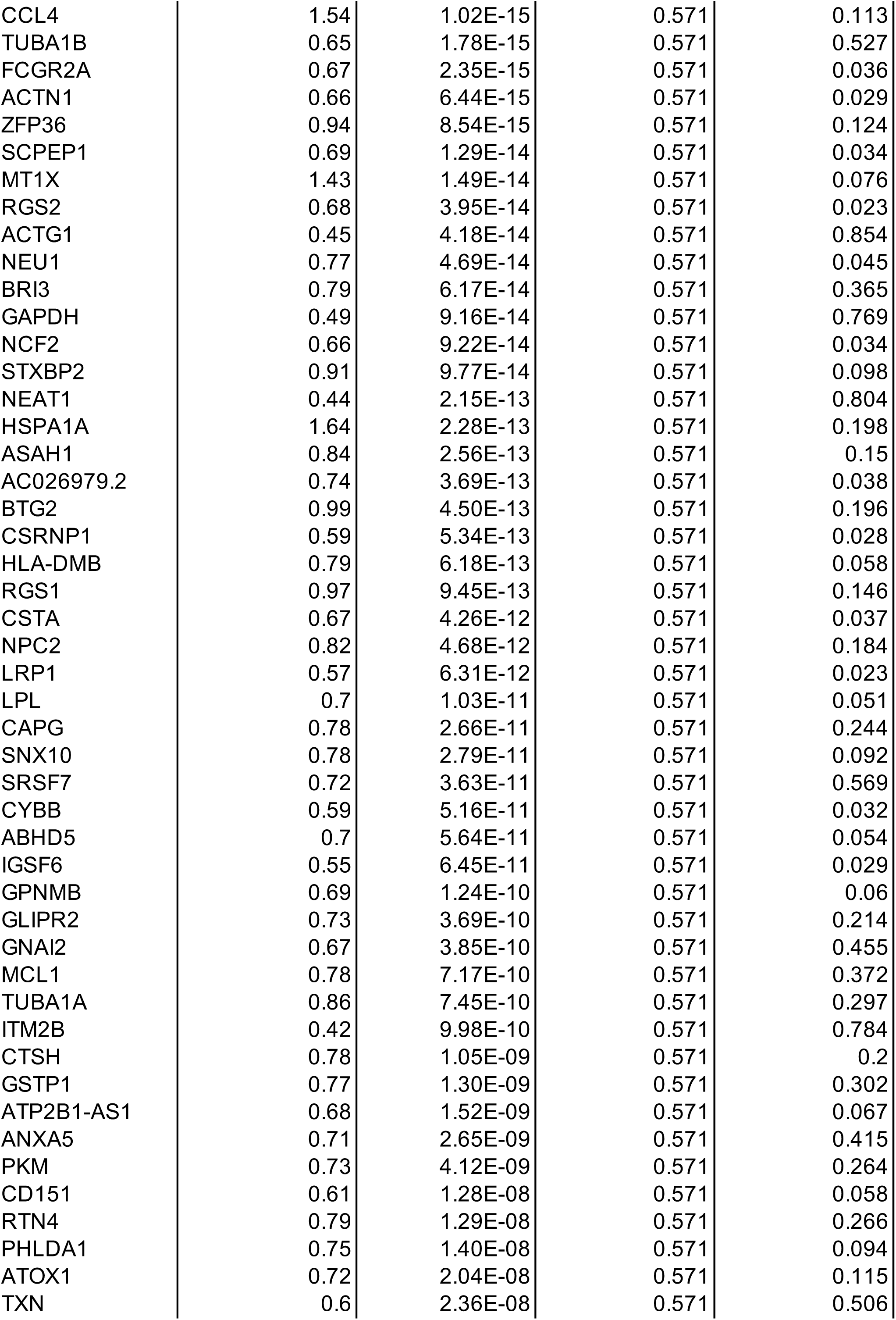
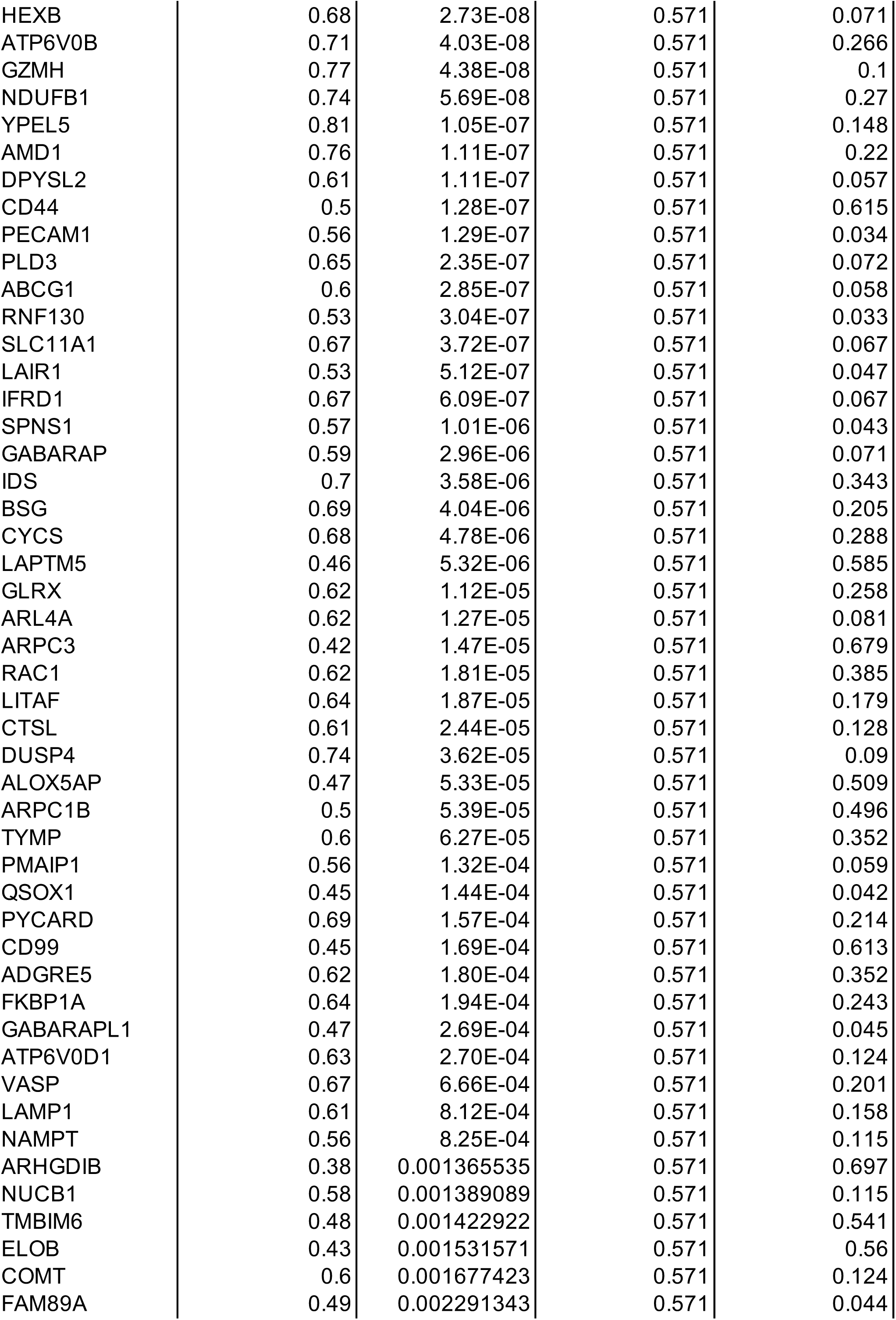
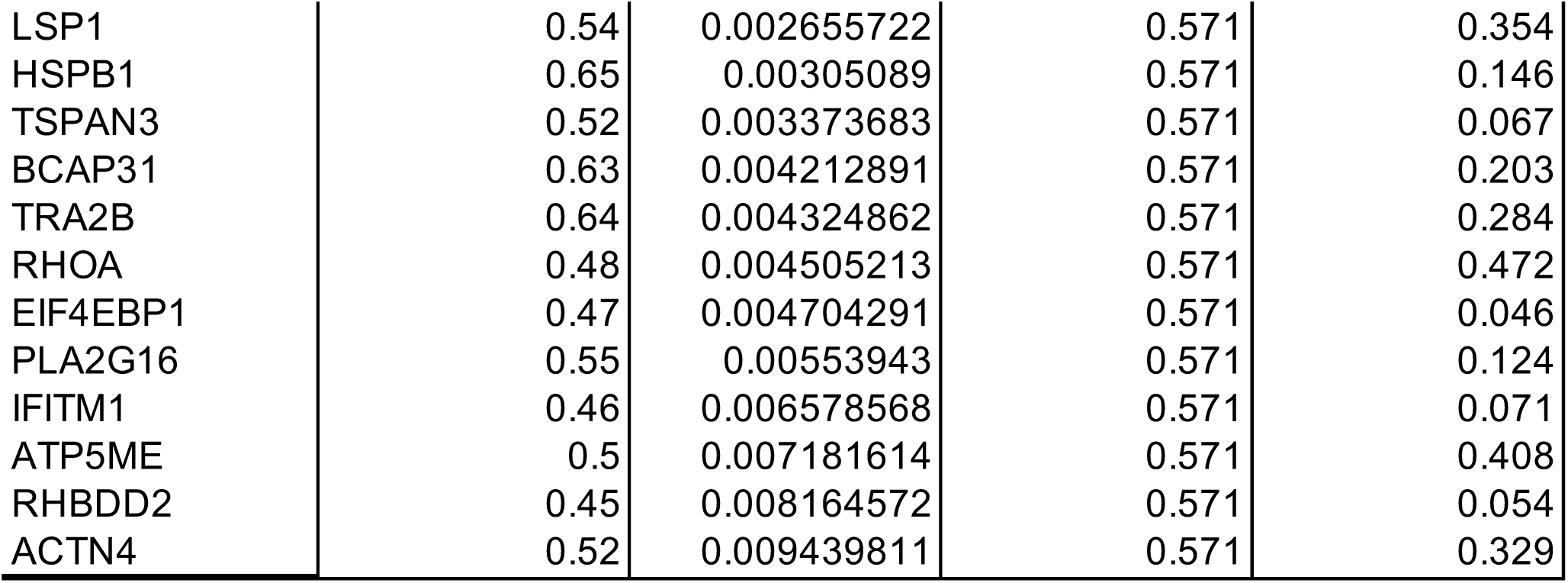
List of Differentially Expressed Genes Upregulated in Disease vs Healthy Subjects.

| Gene | avg_log2FC | p_val_adj | pct.disease | pct.healthy |
|---|---|---|---|---|
| FOS | 3.68 | 6.64E-278 | 0.796 | 0.077 |
| HLA-DRB1 | 2.68 | 9.24E-268 | 0.974 | 0.535 |
| FTH1 | 1.8 | 1.99E-267 | 1 | 0.984 |
| FTL | 1.85 | 1.79E-257 | 1 | 0.997 |
| FABP4 | 2.86 | 6.26E-251 | 0.899 | 0.333 |
| CD74 | 2.29 | 1.22E-246 | 0.972 | 0.626 |
| HLA-DRA | 2.13 | 1.04E-235 | 0.974 | 0.736 |
| APOC1 | 2.39 | 6.12E-219 | 0.92 | 0.524 |
| DUSP1 | 2.95 | 2.29E-211 | 0.754 | 0.148 |
| C1QB | 2.57 | 4.33E-209 | 0.843 | 0.317 |
| C1QA | 2.49 | 3.07E-194 | 0.818 | 0.308 |
| CST3 | 2.49 | 3.26E-193 | 0.743 | 0.179 |
| HLA-DPA1 | 2 | 3.42E-155 | 0.829 | 0.442 |
| IFI27 | 2.14 | 1.19E-140 | 0.641 | 0.163 |
| LYZ | 1.73 | 2.38E-135 | 0.849 | 0.535 |
| KLF6 | 1.63 | 1.96E-133 | 0.924 | 0.718 |
| FOSB | 2.27 | 2.70E-132 | 0.475 | 0.045 |
| ALDH2 | 2.18 | 4.69E-129 | 0.551 | 0.108 |
| SERPINA1 | 2.12 | 4.17E-123 | 0.547 | 0.115 |
| APOE | 1.83 | 3.20E-120 | 0.646 | 0.21 |
| CTSD | 1.81 | 2.10E-118 | 0.758 | 0.424 |
| MARCO | 1.92 | 1.13E-113 | 0.564 | 0.148 |
| HLA-DPB1 | 1.54 | 1.31E-110 | 0.787 | 0.455 |
| GRN | 1.89 | 8.63E-108 | 0.632 | 0.234 |
| JUND | 1.25 | 3.99E-106 | 0.907 | 0.743 |
| TYROBP | 1.79 | 1.61E-105 | 0.617 | 0.217 |
| LGALS3 | 1.35 | 1.86E-102 | 0.85 | 0.6 |
| JUN | 1.55 | 1.63E-100 | 0.905 | 0.629 |
| FCER1G | 1.83 | 3.29E-100 | 0.529 | 0.137 |
| HLA-DQB1 | 1.69 | 2.48E-96 | 0.603 | 0.22 |
| C1QC | 1.74 | 4.02E-88 | 0.454 | 0.102 |
| SERPING1 | 1.7 | 5.46E-87 | 0.395 | 0.062 |
| CES1 | 1.67 | 2.24E-86 | 0.361 | 0.045 |
| TSC22D3 | 1.49 | 2.55E-86 | 0.732 | 0.422 |
| FN1 | 1.72 | 2.09E-85 | 0.422 | 0.084 |
| CRIP1 | 1.52 | 2.10E-81 | 0.639 | 0.284 |
| PPP1R15A | 1.63 | 1.66E-78 | 0.471 | 0.13 |
| S100A6 | 0.8 | 4.39E-74 | 0.975 | 0.948 |
| CCL18 | 1.62 | 6.91E-71 | 0.484 | 0.161 |
| S100A9 | 1.63 | 1.30E-69 | 0.418 | 0.11 |
| HLA-B | 0.54 | 1.43E-67 | 1 | 0.992 |
| PSAP | 1.45 | 3.28E-64 | 0.571 | 0.283 |
| S100A11 | 0.78 | 6.52E-63 | 0.571 | 0.888 |
| CYBA | 0.88 | 2.95E-59 | 0.571 | 0.779 |
| MT2A | 2.21 | 1.22E-56 | 0.571 | 0.149 |
| TSPO | 1.13 | 1.66E-54 | 0.571 | 0.548 |
| S100A8 | 1.34 | 5.92E-54 | 0.571 | 0.047 |
| FCGR3A | 1.4 | 1.18E-53 | 0.571 | 0.063 |
| VIM | 0.64 | 1.65E-53 | 0.571 | 0.941 |
| AC020916.1 | 1.58 | 2.22E-53 | 0.571 | 0.073 |
| CTSS | 1.26 | 3.41E-50 | 0.571 | 0.315 |
| NUPR1 | 1.31 | 4.35E-49 | 0.571 | 0.07 |
| TNFAIP3 | 1.55 | 1.01E-48 | 0.571 | 0.232 |
| HLA-C | 0.56 | 3.11E-48 | 0.571 | 0.958 |
| TREM1 | 1.2 | 3.56E-48 | 0.571 | 0.031 |
| LMNA | 1.22 | 3.84E-48 | 0.571 | 0.124 |
| VSIG4 | 1.24 | 2.03E-44 | 0.571 | 0.051 |
| AIF1 | 1.28 | 4.70E-43 | 0.571 | 0.087 |
| S100A10 | 0.68 | 9.48E-43 | 0.571 | 0.819 |
| CD63 | 0.99 | 1.68E-42 | 0.571 | 0.531 |
| MCEMP1 | 1.14 | 2.45E-41 | 0.571 | 0.06 |
| RHOB | 1.1 | 4.12E-41 | 0.571 | 0.049 |
| JUNB | 1.32 | 5.53E-41 | 0.571 | 0.487 |
| CTSZ | 1.28 | 1.00E-40 | 0.571 | 0.185 |
| FBP1 | 1.21 | 3.56E-39 | 0.571 | 0.1 |
| S100A4 | 0.69 | 1.63E-38 | 0.571 | 0.859 |
| CD81 | 1.12 | 3.00E-38 | 0.571 | 0.416 |
| UBC | 0.71 | 5.32E-38 | 0.571 | 0.861 |
| MS4A7 | 1.2 | 4.76E-37 | 0.571 | 0.106 |
| CXCL16 | 1.11 | 1.66E-36 | 0.571 | 0.049 |
| ACP5 | 1.27 | 1.43E-35 | 0.571 | 0.146 |
| MYL6 | 0.66 | 4.65E-35 | 0.571 | 0.831 |
| UBB | 0.78 | 8.19E-34 | 0.571 | 0.677 |
| OAZ1 | 0.75 | 1.37E-33 | 0.571 | 0.764 |
| ANXA2 | 0.94 | 2.19E-32 | 0.571 | 0.53 |
| NFKBIA | 1.08 | 3.22E-32 | 0.571 | 0.225 |
| CTSB | 1.08 | 4.08E-32 | 0.571 | 0.275 |
| PLBD1 | 0.94 | 5.08E-32 | 0.571 | 0.021 |
| FCGRT | 1.17 | 5.88E-32 | 0.571 | 0.097 |
| LGALS3BP | 1.14 | 2.78E-31 | 0.571 | 0.09 |
| OLR1 | 1.02 | 2.83E-31 | 0.571 | 0.047 |
| CSTB | 0.96 | 2.90E-30 | 0.571 | 0.472 |
| HP | 0.8 | 3.10E-29 | 0.571 | 0.011 |
| CFD | 1.1 | 4.16E-29 | 0.571 | 0.094 |
| SCD | 1.08 | 7.21E-29 | 0.571 | 0.084 |
| SRGN | 0.8 | 7.67E-29 | 0.571 | 0.578 |
| SPI1 | 0.99 | 1.05E-28 | 0.571 | 0.049 |
| CD9 | 1.02 | 3.32E-28 | 0.571 | 0.099 |
| TAGLN2 | 0.86 | 4.53E-28 | 0.571 | 0.484 |
| GCHFR | 1.01 | 2.00E-27 | 0.571 | 0.187 |
| LY6E | 0.8 | 3.47E-27 | 0.571 | 0.503 |
| CD68 | 1.03 | 5.33E-27 | 0.571 | 0.08 |
| COTL1 | 0.9 | 1.26E-26 | 0.571 | 0.473 |
| CD69 | 1.13 | 1.42E-26 | 0.571 | 0.319 |
| SAT1 | 0.8 | 3.01E-26 | 0.571 | 0.43 |
| RGCC | 0.99 | 4.50E-26 | 0.571 | 0.291 |
| HLA-DQA1 | 0.98 | 6.47E-26 | 0.571 | 0.141 |
| LST1 | 1.04 | 8.14E-26 | 0.571 | 0.128 |
| IFNG | 1.55 | 1.07E-25 | 0.571 | 0.111 |
| MSR1 | 0.94 | 2.82E-25 | 0.571 | 0.054 |
| NR4A2 | 0.79 | 7.27E-25 | 0.571 | 0.011 |
| SH3BGRL3 | 0.55 | 1.09E-24 | 0.571 | 0.887 |
| MRC1 | 0.91 | 3.66E-24 | 0.571 | 0.073 |
| HLA-DMA | 1.04 | 8.47E-24 | 0.571 | 0.126 |
| LTA4H | 0.94 | 9.61E-24 | 0.571 | 0.095 |
| LGALS1 | 0.7 | 1.99E-23 | 0.571 | 0.582 |
| HSPA1B | 1.31 | 3.83E-23 | 0.571 | 0.036 |
| CEBPB | 0.92 | 4.76E-23 | 0.571 | 0.367 |
| CXCL3 | 0.66 | 5.38E-23 | 0.571 | 0.008 |
| RETN | 1.01 | 5.58E-23 | 0.571 | 0.076 |
| PHLDA3 | 0.83 | 2.24E-22 | 0.571 | 0.027 |
| CYP27A1 | 0.75 | 5.05E-22 | 0.571 | 0.03 |
| PLAUR | 0.84 | 6.64E-22 | 0.571 | 0.033 |
| MYADM | 0.95 | 1.87E-21 | 0.571 | 0.197 |
| IGFBP2 | 0.73 | 3.14E-21 | 0.571 | 0.021 |
| GLUL | 0.98 | 6.55E-21 | 0.571 | 0.182 |
| TIMP2 | 0.87 | 1.08E-20 | 0.571 | 0.046 |
| S100A13 | 0.67 | 2.03E-20 | 0.571 | 0.016 |
| ALOX5 | 0.86 | 2.90E-20 | 0.571 | 0.061 |
| C1orf162 | 0.91 | 3.11E-20 | 0.571 | 0.161 |
| PDLIM1 | 0.76 | 1.64E-19 | 0.571 | 0.027 |
| AC044849.1 | 0.83 | 2.72E-19 | 0.571 | 0.027 |
| MNDA | 0.81 | 7.00E-19 | 0.571 | 0.034 |
| TMSB10 | 0.36 | 9.42E-19 | 0.571 | 0.968 |
| CTSC | 0.71 | 1.24E-18 | 0.571 | 0.475 |
| CD52 | 0.48 | 1.71E-18 | 0.571 | 0.854 |
| RBP4 | 0.72 | 1.83E-18 | 0.571 | 0.036 |
| CFL1 | 0.46 | 2.44E-18 | 0.571 | 0.826 |
| PFN1 | 0.53 | 2.91E-18 | 0.571 | 0.762 |
| DDX5 | 0.55 | 7.52E-18 | 0.571 | 0.821 |
| SERF2 | 0.49 | 8.87E-18 | 0.571 | 0.854 |
| FABP5 | 0.83 | 2.52E-17 | 0.571 | 0.24 |
| HSP90AA1 | 0.7 | 2.72E-17 | 0.571 | 0.788 |
| MGST3 | 0.82 | 3.02E-17 | 0.571 | 0.367 |
| INHBA | 0.79 | 3.44E-17 | 0.571 | 0.038 |
| ANXA1 | 0.56 | 6.64E-17 | 0.571 | 0.677 |
| MS4A4A | 0.69 | 1.02E-16 | 0.571 | 0.029 |
| PRDX1 | 0.77 | 1.41E-16 | 0.571 | 0.464 |
| GAA | 0.68 | 4.50E-16 | 0.571 | 0.023 |
| DNAJB1 | 1.2 | 9.33E-16 | 0.571 | 0.096 |
| SDCBP | 0.93 | 9.80E-16 | 0.571 | 0.188 |
| CCL4 | 1.54 | 1.02E-15 | 0.571 | 0.113 |
| TUBA1B | 0.65 | 1.78E-15 | 0.571 | 0.527 |
| FCGR2A | 0.67 | 2.35E-15 | 0.571 | 0.036 |
| ACTN1 | 0.66 | 6.44E-15 | 0.571 | 0.029 |
| ZFP36 | 0.94 | 8.54E-15 | 0.571 | 0.124 |
| SCPEP1 | 0.69 | 1.29E-14 | 0.571 | 0.034 |
| MT1X | 1.43 | 1.49E-14 | 0.571 | 0.076 |
| RGS2 | 0.68 | 3.95E-14 | 0.571 | 0.023 |
| ACTG1 | 0.45 | 4.18E-14 | 0.571 | 0.854 |
| NEU1 | 0.77 | 4.69E-14 | 0.571 | 0.045 |
| BRI3 | 0.79 | 6.17E-14 | 0.571 | 0.365 |
| GAPDH | 0.49 | 9.16E-14 | 0.571 | 0.769 |
| NCF2 | 0.66 | 9.22E-14 | 0.571 | 0.034 |
| STXBP2 | 0.91 | 9.77E-14 | 0.571 | 0.098 |
| NEAT1 | 0.44 | 2.15E-13 | 0.571 | 0.804 |
| HSPA1A | 1.64 | 2.28E-13 | 0.571 | 0.198 |
| ASAH1 | 0.84 | 2.56E-13 | 0.571 | 0.15 |
| AC026979.2 | 0.74 | 3.69E-13 | 0.571 | 0.038 |
| BTG2 | 0.99 | 4.50E-13 | 0.571 | 0.196 |
| CSRNP1 | 0.59 | 5.34E-13 | 0.571 | 0.028 |
| HLA-DMB | 0.79 | 6.18E-13 | 0.571 | 0.058 |
| RGS1 | 0.97 | 9.45E-13 | 0.571 | 0.146 |
| CSTA | 0.67 | 4.26E-12 | 0.571 | 0.037 |
| NPC2 | 0.82 | 4.68E-12 | 0.571 | 0.184 |
| LRP1 | 0.57 | 6.31E-12 | 0.571 | 0.023 |
| LPL | 0.7 | 1.03E-11 | 0.571 | 0.051 |
| CAPG | 0.78 | 2.66E-11 | 0.571 | 0.244 |
| SNX10 | 0.78 | 2.79E-11 | 0.571 | 0.092 |
| SRSF7 | 0.72 | 3.63E-11 | 0.571 | 0.569 |
| CYBB | 0.59 | 5.16E-11 | 0.571 | 0.032 |
| ABHD5 | 0.7 | 5.64E-11 | 0.571 | 0.054 |
| IGSF6 | 0.55 | 6.45E-11 | 0.571 | 0.029 |
| GPNMB | 0.69 | 1.24E-10 | 0.571 | 0.06 |
| GLIPR2 | 0.73 | 3.69E-10 | 0.571 | 0.214 |
| GNAI2 | 0.67 | 3.85E-10 | 0.571 | 0.455 |
| MCL1 | 0.78 | 7.17E-10 | 0.571 | 0.372 |
| TUBA1A | 0.86 | 7.45E-10 | 0.571 | 0.297 |
| ITM2B | 0.42 | 9.98E-10 | 0.571 | 0.784 |
| CTSH | 0.78 | 1.05E-09 | 0.571 | 0.2 |
| GSTP1 | 0.77 | 1.30E-09 | 0.571 | 0.302 |
| ATP2B1-AS1 | 0.68 | 1.52E-09 | 0.571 | 0.067 |
| ANXA5 | 0.71 | 2.65E-09 | 0.571 | 0.415 |
| PKM | 0.73 | 4.12E-09 | 0.571 | 0.264 |
| CD151 | 0.61 | 1.28E-08 | 0.571 | 0.058 |
| RTN4 | 0.79 | 1.29E-08 | 0.571 | 0.266 |
| PHLDA1 | 0.75 | 1.40E-08 | 0.571 | 0.094 |
| ATOX1 | 0.72 | 2.04E-08 | 0.571 | 0.115 |
| TXN | 0.6 | 2.36E-08 | 0.571 | 0.506 |
| HEXB | 0.68 | 2.73E-08 | 0.571 | 0.071 |
| ATP6V0B | 0.71 | 4.03E-08 | 0.571 | 0.266 |
| GZMH | 0.77 | 4.38E-08 | 0.571 | 0.1 |
| NDUFB1 | 0.74 | 5.69E-08 | 0.571 | 0.27 |
| YPEL5 | 0.81 | 1.05E-07 | 0.571 | 0.148 |
| AMD1 | 0.76 | 1.11E-07 | 0.571 | 0.22 |
| DPYSL2 | 0.61 | 1.11E-07 | 0.571 | 0.057 |
| CD44 | 0.5 | 1.28E-07 | 0.571 | 0.615 |
| PECAM1 | 0.56 | 1.29E-07 | 0.571 | 0.034 |
| PLD3 | 0.65 | 2.35E-07 | 0.571 | 0.072 |
| ABCG1 | 0.6 | 2.85E-07 | 0.571 | 0.058 |
| RNF130 | 0.53 | 3.04E-07 | 0.571 | 0.033 |
| SLC11A1 | 0.67 | 3.72E-07 | 0.571 | 0.067 |
| LAIR1 | 0.53 | 5.12E-07 | 0.571 | 0.047 |
| IFRD1 | 0.67 | 6.09E-07 | 0.571 | 0.067 |
| SPNS1 | 0.57 | 1.01E-06 | 0.571 | 0.043 |
| GABARAP | 0.59 | 2.96E-06 | 0.571 | 0.071 |
| IDS | 0.7 | 3.58E-06 | 0.571 | 0.343 |
| BSG | 0.69 | 4.04E-06 | 0.571 | 0.205 |
| CYCS | 0.68 | 4.78E-06 | 0.571 | 0.288 |
| LAPTM5 | 0.46 | 5.32E-06 | 0.571 | 0.585 |
| GLRX | 0.62 | 1.12E-05 | 0.571 | 0.258 |
| ARL4A | 0.62 | 1.27E-05 | 0.571 | 0.081 |
| ARPC3 | 0.42 | 1.47E-05 | 0.571 | 0.679 |
| RAC1 | 0.62 | 1.81E-05 | 0.571 | 0.385 |
| LITAF | 0.64 | 1.87E-05 | 0.571 | 0.179 |
| CTSL | 0.61 | 2.44E-05 | 0.571 | 0.128 |
| DUSP4 | 0.74 | 3.62E-05 | 0.571 | 0.09 |
| ALOX5AP | 0.47 | 5.33E-05 | 0.571 | 0.509 |
| ARPC1B | 0.5 | 5.39E-05 | 0.571 | 0.496 |
| TYMP | 0.6 | 6.27E-05 | 0.571 | 0.352 |
| PMAIP1 | 0.56 | 1.32E-04 | 0.571 | 0.059 |
| QSOX1 | 0.45 | 1.44E-04 | 0.571 | 0.042 |
| PYCARD | 0.69 | 1.57E-04 | 0.571 | 0.214 |
| CD99 | 0.45 | 1.69E-04 | 0.571 | 0.613 |
| ADGRE5 | 0.62 | 1.80E-04 | 0.571 | 0.352 |
| FKBP1A | 0.64 | 1.94E-04 | 0.571 | 0.243 |
| GABARAPL1 | 0.47 | 2.69E-04 | 0.571 | 0.045 |
| ATP6V0D1 | 0.63 | 2.70E-04 | 0.571 | 0.124 |
| VASP | 0.67 | 6.66E-04 | 0.571 | 0.201 |
| LAMP1 | 0.61 | 8.12E-04 | 0.571 | 0.158 |
| NAMPT | 0.56 | 8.25E-04 | 0.571 | 0.115 |
| ARHGDIB | 0.38 | 0.001365535 | 0.571 | 0.697 |
| NUCB1 | 0.58 | 0.001389089 | 0.571 | 0.115 |
| TMBIM6 | 0.48 | 0.001422922 | 0.571 | 0.541 |
| ELOB | 0.43 | 0.001531571 | 0.571 | 0.56 |
| COMT | 0.6 | 0.001677423 | 0.571 | 0.124 |
| FAM89A | 0.49 | 0.002291343 | 0.571 | 0.044 |
| LSP1 | 0.54 | 0.002655722 | 0.571 | 0.354 |
| HSPB1 | 0.65 | 0.00305089 | 0.571 | 0.146 |
| TSPAN3 | 0.52 | 0.003373683 | 0.571 | 0.067 |
| BCAP31 | 0.63 | 0.004212891 | 0.571 | 0.203 |
| TRA2B | 0.64 | 0.004324862 | 0.571 | 0.284 |
| RHOA | 0.48 | 0.004505213 | 0.571 | 0.472 |
| EIF4EBP1 | 0.47 | 0.004704291 | 0.571 | 0.046 |
| PLA2G16 | 0.55 | 0.00553943 | 0.571 | 0.124 |
| IFITM1 | 0.46 | 0.006578568 | 0.571 | 0.071 |
| ATP5ME | 0.5 | 0.007181614 | 0.571 | 0.408 |
| RHBDD2 | 0.45 | 0.008164572 | 0.571 | 0.054 |
| ACTN4 | 0.52 | 0.009439811 | 0.571 | 0.329 |

Taken together, our study suggests that exposure to Be results in enhanced activation of CD4^+^ T cells that contribute to inflammation and perpetuation of disease. Single-cell sequencing revealed significantly increased expression of *CD74* and *FABP4* (adjusted p < 0.01), genes predominantly used to define resident alveolar macrophages (15). Based on recent reports, fatty-acid binding proteins (FABPs) are upregulated in CD4^+^ T cells and regulate multiple functions (16). Therefore, further studies assessing the function of these molecules in CD4^+^ T cells in granulomatous lung diseases using human samples and mouse models will help identify new targets for drug discovery or immunotherapy, as well as define pathways specific to pathogenic CD4^+^ T cells in beryllium disease.

## Methods

### Single-cell sequencing

Bronchoalveolar lavage samples were collected from two Be-exposed subjects at National Jewish Health (NJH) Granuloma Clinic. Subjects provided informed consent prior to inclusion in the study, and the protocol was approved by the Human Subject Institutional Review Boards at the University of Colorado Anschutz Medical Campus and National Jewish Health. Dataset for healthy control samples was obtained from Gene Expression Omnibus, accession code GSE151928 (17). Cells were thawed and rested in culture overnight in complete medium. Single cells were flow-sorted using propidium iodide to exclude dead cells. The sorted cells were resuspended in PBS buffer, recounted, and processed for the 10X Genomics 3’v2 Kit as specified by the manufacturer’s protocol for each patient sample. Approximately 5000 cells were loaded in the wells of a 10X Genomics Chromium controller to create GEM emulsions, targeting a recovery of 3000 cells. Reverse transcription, cDNA amplification, and RNA-sequencing library preparation were performed using the GEM emulsion. Libraries were sequenced on the Illumina Hi-Seq 4000 platform, using 75 bp paired-end reads.

### Single-cell sequencing data and GO pathway enrichment analysis

After Cell Ranger (10x Genomics) processing, scRNA-seq analyses were carried out with the tool suite Seurat (6), utilizing cell pairwise correspondence dataset integration, UMAP dimension reduction, and SNN clustering. Differential gene expression was defined by a Wilcoxon signed-rank test, with thresholds of adjusted p-value ≤ 0.01 and log2 fold change of over 0.25. The top 200 differentially expressed genes, ranked by adjusted p-value, were used for GO term enrichment analysis.

## Acknowledgement

This work was supported by National Institutes of Health grant (HL152756), TRSP, FSR to S.M.A.

## Notes

https://www.ncbi.nlm.nih.gov/geo/query/acc.cgi?acc=GSE181174

https://www.ncbi.nlm.nih.gov/geo/query/acc.cgi?acc=GSE325899

### Competing Interest Statement

The authors have declared no competing interest.

## References

1. Fontenot AP, Maier LA. Genetic susceptibility and immune-mediated destruction in beryllium-induced disease. Trends Immunol 2005; 26: 543–549.

2. McKee AS, Fontenot AP. Interplay of innate and adaptive immunity in metal-induced hypersensitivity. Curr Opin Immunol 2016; 42: 25–30.

3. Saltini C, Winestock K, Kirby M, Pinkston P, Crystal RG. Maintenance of alveolitis in patients with chronic beryllium disease by beryllium-specific helper T cells. N Engl J Med 1989; 320: 1103–1109.

4. Newman LS. Significance of the blood beryllium lymphocyte proliferation test. Environmental Health Perspectives 1996; 104 Suppl 5: 953–956.

5. Mroz MM, Kreiss K, Lezotte DC, Campbell PA, Newman LS. Reexamination of the blood lymphocyte transformation test in the diagnosis of chronic beryllium disease. J Allergy Clin Immunol 1991; 88: 54–60.

6. Stuart T, Butler A, Hoffman P, Hafemeister C, Papalexi E, Mauck WM, 3rd, Hao Y, Stoeckius M, Smibert P, Satija R. Comprehensive Integration of Single-Cell Data. Cell 2019; 177: 1888–1902 e1821.

7. Travaglini KJ, Nabhan AN, Penland L, Sinha R, Gillich A, Sit RV, Chang S, Conley SD, Mori Y, Seita J, Berry GJ, Shrager JB, Metzger RJ, Kuo CS, Neff N, Weissman IL, Quake SR, Krasnow MA. A molecular cell atlas of the human lung from single-cell RNA sequencing. Nature 2020; 587: 619–625.

8. McInnes L, Healy J, Melville J. UMAP: Uniform manifold approximation and projection for dimension reduction. Stat Mach Learn 2018; arXiv:1802.03426.

9. Fontenot AP, Canavera SJ, Gharavi L, Newman LS, Kotzin BL. Target organ localization of memory CD4^+^ T cells in patients with chronic beryllium disease. J Clin Invest 2002; 110: 1473–1482.

10. Frizzell H, Fonseca R, Christo SN, Evrard M, Cruz-Gomez S, Zanluqui NG, von Scheidt B, Freestone D, Park SL, McWilliam HEG, Villadangos JA, Carbone FR, Mackay LK. Organ-specific isoform selection of fatty acid-binding proteins in tissue-resident lymphocytes. Sci Immunol 2020; 5.

11. Hu H, Nau M, Ehrenberg P, Chenine AL, Macedo C, Zhou Y, Daye ZJ, Wei Z, Vahey M, Michael NL, Kim JH, Marovich M, Ratto-Kim S. Distinct gene-expression profiles associated with the susceptibility of pathogen-specific CD4 T cells to HIV-1 infection. Blood 2013; 121: 1136–1144.

12. Khamashta M, Merrill JT, Werth VP, Furie R, Kalunian K, Illei GG, Drappa J, Wang L, Greth W, investigators CDs. Sifalimumab, an anti-interferon-alpha monoclonal antibody, in moderate to severe systemic lupus erythematosus: a randomised, double-blind, placebo-controlled study. Ann Rheum Dis 2016; 75: 1909–1916.

13. Street K, Risso D, Fletcher RB, Das D, Ngai J, Yosef N, Purdom E, Dudoit S. Slingshot: cell lineage and pseudotime inference for single-cell transcriptomics. BMC Genomics 2018; 19: 477.

14. Falta MT, Raza M, Nevienski CJ, Brunetti TM, Fu R, Tucker RM, Gaballa JM, Minhajuddin F, Alula KM, Dinarello A, Mack DG, Martin AK, Onyiah JC, Yarnell M, Francis P, Fry TJ, Maier LA, Fontenot AP, Dinarello CA, Atif SM. Reduced CCL/Be-specific CD4+ T cells in CCL3-deficient or peptide-MHCII CAR-T cell-treated mice. JCI Insight 2026; 11.

15. Liao SY, Atif SM, Mould K, Konigsberg IR, Fu R, Davidson E, Li L, Fontenot AP, Maier LA, Yang IV. Single-cell RNA sequencing identifies macrophage transcriptional heterogeneities in granulomatous diseases. Eur Respir J 2021.

16. Field CS, Baixauli F, Kyle RL, Puleston DJ, Cameron AM, Sanin DE, Hippen KL, Loschi M, Thangavelu G, Corrado M, Edwards-Hicks J, Grzes KM, Pearce EJ, Blazar BR, Pearce EL. Mitochondrial Integrity Regulated by Lipid Metabolism Is a Cell-Intrinsic Checkpoint for Treg Suppressive Function. Cell Metab 2020; 31: 422–437 e425.

17. Mould KJ, Moore CM, McManus SA, McCubbrey AL, McClendon JD, Griesmer CL, Henson PM, Janssen WJ. Airspace Macrophages and Monocytes Exist in Transcriptionally Distinct Subsets in Healthy Adults. Am J Respir Crit Care Med 2021; 203: 946–956.

